# CRISPR-based Genome Editing of a Diurnal Rodent, Nile Grass Rat (*Arvicanthis niloticus)*

**DOI:** 10.1101/2023.08.23.553600

**Authors:** Huirong Xie, Katrina Linning-Duffy, Elena Y. Demireva, Huishi Toh, Bana Abolibdeh, Jiaming Shi, Bo Zhou, Shigeki Iwase, Lily Yan

**Affiliations:** Transgenic and Genome Editing Facility, Institute for Quantitative Health Science & Engineering, Research Technology Support Facility, Michigan State University; Department of Psychology, Michigan State University; Neuroscience Research Institute, University of California Santa Barbara; Department of Human Genetics, University of Michigan Medical School; Department of Pediatrics, University of Michigan Medical School; Neuroscience Program, Michigan State University

**Keywords:** Nile Grass Rat, CRISPR-Cas9, Genome Editing, Rai1 KO, Diurnal Rodent

## Abstract

Diurnal and nocturnal mammals have evolved distinct pathways to optimize survival for their chronotype-specific lifestyles. Conventional rodent models, being nocturnal, may not sufficiently recapitulate the biology of diurnal humans in health and disease. Although diurnal rodents are potentially advantageous for translational research, until recently, they have not been genetically tractable. Here, we address this major limitation by demonstrating the first successful CRISPR genome editing of the Nile grass rat (*Arvicanthis niloticus*), a valuable diurnal rodent. We establish methods for superovulation; embryo development, manipulation, and culture; and pregnancy maintenance to guide future genome editing of this and other diurnal rodent species.

## Background

Model organisms are essential for biomedical research in understanding physiology and pathology relevant to human health and disease. Commonly used animal models in biomedical research including laboratory mice or rats are nocturnal (night-active), while humans are diurnal (day-active). Diurnal and nocturnal mammals have acquired different adaptations through the evolution of numerous pathways to optimize survival for a day- or night-active lifestyle[1]. An internal time-keeping system, namely the circadian clock system, has evolved to predict and prepare animals for the daily fluctuations in their environment. Circadian systems coordinate the temporal organizations of molecular, cellular, and physiological processes across the body to ensure the functions of cells, tissues, and organs are synchronized with the environmental day-night cycle[2]. In mammals, this system is organized in a hierarchical manner, with the principal brain clock within the hypothalamic suprachiasmatic nucleus (SCN), coordinating the circadian rhythms of subordinate clocks in other brain regions and in peripheral tissues and organs[3]. The expression of core circadian genes within the SCN shows the same temporal dynamics i.e., peaking at the same time in diurnal and nocturnal animals, however, other brain regions and peripheral organs show complex differences between the two chronotypes. Large-scale transcriptomic studies revealed that core circadian genes’ peak expression shifted by 6-15 hours between nocturnal (mouse) and diurnal (baboon) species depending on tissue types[4, 5]. Therefore, the circadian system in nocturnal and diurnal species differs in a more complex manner than a simply inverted daily pattern[1], which likely involves distinct wiring of neural circuits and gene-regulatory networks[4-7]. Furthermore, evidence suggests that experimentation during nocturnal rodents’ inactive phase can be a major cause of human clinical trial failures of drug candidates proven to be effective in preclinical mouse models[8]. For these reasons, diurnal rodents are advantageous over nocturnal ones for translational research[9]. A major limitation of diurnal rodents in biomedical research is that they have not been genetically tractable. The present study aimed to overcome this barrier and develop methods for gene editing in a diurnal rodent, the Nile grass rat (*Arvicanthis niloticus)*.

Nile grass rats, together with laboratory mice (*Mus musculus*) and laboratory rats (*Rattus norvegicus*), are members of the family Muridae[10], and these species are likely to have diverged from a common ancestor relatively recently[1]. Nile grass rats, like mice, attain reproductive maturity rapidly, have a 24-day gestation period and mate on postpartum estrus, which makes maintenance of a colony relatively simple[11]. Nile grass rats are clearly diurnal both in nature and in the laboratory, as indicated by their patterns of activity, sleep, mating behavior, body temperature and secretion of luteinizing hormones[1]. Their retinal anatomy and retinorecipient brain regions are also typical for animals active during the daytime[12-14]. The Nile grass rat colony at Michigan State University was established in 1993 from a cohort of animals captured from the Maasai Mara National Reserve in Kenya[15]. The colony has been maintained since then; and animals derived from this colony have been shared with numerous research groups that investigate circadian rhythms and sleep, affective behaviors, cognitive function, immune function, metabolic syndromes, ophthalmology, and evolutionary biology. Despite being a well-established diurnal rodent model, Nile grass rats have not been genetically tractable because their complete genome sequence and an established genome editing protocol have not been available. Recently, the Vertebrate Genome Project[16, 17] released the initial build of the Nile grass rat genome[18], opening up an opportunity for genome editing in this species.

In addition to a sequenced genome, another requirement for gene targeting in a specific organism is the availability of gene editing technologies. During the last several decades, precise gene editing technology in mice and rats has progressed from the time-consuming and costly embryonic stem cell-based targeting[19-21], to rapid genome targeting approaches utilizing zinc finger nucleases (ZFN)[22-24] and transcription activator-like effector nucleases (TALEN)[25-28]. In 2013, CRISPR-Cas9 was first used to generate precise deletions and point mutations of two genes, *Tet1* and *Tet2* at once by microinjecting mRNA of Cas9 nuclease and guide RNAs into mouse zygotes[29]. Since then, the CRISPR-Cas9 technology has been applied broadly in creating genome modified models in many different species[30-35]. Furthermore, delivery methods also expanded beyond microinjection, with electroporation of Cas9 mRNA or protein and gRNA into zygotes becoming an efficient genome editing approach[36-38]. The improved Genome-editing via Oviductal Nucleic Acids Delivery (i-GONAD) method has further enabled the in vivo delivery of CRISPR components without the need for embryo culture or pseudopregnant recipients[39-45].

In the present work, taking advantage of CRISPR-Cas9 and i-GONAD, we developed a method for genome editing of the Nile grass rat. To our knowledge, this study demonstrates genome editing of this well-established diurnal rodent model for the first time. The first targeted gene in this species is the Retinoic acid-induced 1 (*Rai1*) gene, whose haploinsufficiency is responsible for Smith-Magenis Syndrome, a neurodevelopmental disorder characterized by sleep disturbance in humans. We also succeeded in several critical steps required for gene targeting, including superovulation and embryo culture, which will allow for direct in vitro embryonic manipulation (microinjection and electroporation) of a variety of genome editing reagents beyond CRISPR-Cas9, thereby paving the way for future efforts to equip this diurnal model with a variety of molecular and genetic tools currently available for conventional laboratory mice or rats.

## Results

### Superovulation of Nile grass rats

In order to produce a high number of fertilized grass rat embryos for genome editing, we attempted to establish a superovulation protocol by varying the timing of hormone treatment and egg collection as outlined below. The egg yield and fertilization rate were then compared to those from a natural mating cohort.

### Timing of hormone treatment and egg collection

Due to the lack of knowledge about the reproductive biology of this species, we designed superovulation protocols based on observations from grass rat breeding and standard superovulation protocols in mouse and rat. Our colony breeding records showed that a notable number of first litters were born between 26 and 30 days after males and females were paired, indicating that day 3 and day 4 post pairing is likely the early receptive mating window. Therefore, superovulation protocols were set to administer hCG on day 3 or 4 after PMSG priming. Embryo yields from females that underwent superovulation with PMSG and hCG were compared to those from unassisted natural mating. Collectively 6 out of 8 groups (Table 1, group # 3-8) of superovulated females produced 20 eggs per female on average (mean ± SEM: 20.8 ± 2.2), significantly higher than the number of eggs (5 ± 0.9) produced by natural mating (t-test, t_31_ = 4.88, p < 0.001). In those 6 groups (#3-8), PMSG was administered between 6:00 am and 11:00 am (day 1), hCG was administered 48 to 57 hours later between 2:00 pm and 4:00 pm (day 3), and eggs were collected on day 4 at 9:00 am or 5:00 pm (19 to 27 hours post-hCG, group #4, 5, 8) to time the embryo development at pronuclear stage, or day 5 at 9:00 am, 10:00 am or 2:00 pm (40 to 51 hours post-hCG, group #3, 6, 7) with the aim to obtain 2-cell staged embryos. A shorter PMSG-hCG interval (36 to 48 hours) and an earlier egg collection (on day 3) was tested in groups #1 and #2 which resulted in a lower yield of eggs (4.6 ± 1.1), comparable to that from the natural mating group (t-test, t_18_ = 0.32, p = 0.75). While the dosage of PMSG and hCG was kept constant (15 IU each, ∼150 IU/kg) except in group #6 (20 IU each), eight combinations of different timing and intervals of PMSG and hCG administration were tested, six of them (group #3-8) resulted in higher egg yield than natural mating. In group #7, the yield reached nearly 30 eggs per female, 5-fold higher compared to natural mating (Table 1). In summary, superovulation of grass rat females can be achieved, and the current protocol is sufficient to produce large number of oocytes.

**Table 1:** Egg and embryo yields from superovulated and natural mating females.

| **Group**<br>**(sample**<br>**size)** | **PMSG**<br>**(day 1)**<br>**(+/- male)** | **PMSG-hCG**<br>**interval (day, time**<br>**of hCG, +/- male)** | **hCG-harvest**<br>**interval**<br>**(day, time)** | **# sperm (+)**<br>**females (eggs)**<br>**harvested** | **# sperm (-)**<br>**females (eggs)**<br>**harvested** | **# fertilized eggs**<br>**(fertilization**<br>**rate<sup>1</sup>)** | **# eggs/**<br>**female** |
| --- | --- | --- | --- | --- | --- | --- | --- |
| #1<br>(n = 3) | 6 am | 36 h, (day2, 6pm),<br>+ male | 24 h<br>(day3, 6pm) | 1 (2) | 2 (4) | 0 | 3 |
| #2<br>(n = 8) | 6 am | 48 h, (day3, 6am),<br>+ male | 12 h<br>(day3, 6pm) | 1 (11) | 5 (24) | 3 (27.3%) | 6 |
| #3<br>(n = 3) | 6 am | 48 h, (day3, 6am),<br>+ male | 51 h<br>(day5, 9am) | 1 (11) | 2 (45) | 11 (100%) | 18.7 |
| #4<br>(n = 3) | 6 am | 56 h, (day3, 2pm),<br>+ male | 19 h<br>(day4, 9am) | 2 (29) | 1 (27) | 0 (0) | 18.7 |
| #5<br>(n = 2) | 6 am | 56 h, (day3, 2pm),<br>+ male | 27 h<br>(day4, 5pm) | 1 (13) | 0 | 5 (38.5%) | 13 |
| #6<br>(n = 7) | 7 am<br>+ male | 57 h, (day3, 4pm) | 46 h<br>(day5, 2pm) | 0 | 7 (132) | 0 (0) | 18.9 |
| #7<br>(n = 5) | 10 am<br>+ male | 56 h, (day3, 6pm) | 40 h<br>(day5, 10am) | 0 | 5 (163) | 0 (0) | 32.6 |
| #8<br>(n = 6) | 11 am<br>+ male | 51 h, (day3, 2pm) | 27 h<br>(day4, 5pm) | 3 (44) | 0 | 5 (11.6%) | 14.7 |
| Natural<br>mating<br>(n = 17) | No PMSG,<br>+ male | No hCG | 100 h post-pairing (day 5) | 11 (55) | 0 | 51 (92.7%) | 5 |

PMSG/hCG were administered at 15 IU in all superovulation groups except group #6, which received 20 IU.
<sup>1</sup>Fertilization rates were determined as the number of fertilized eggs divided by the total number of eggs from sperm positive females for each group of females.

### Fertilization rate

Although the number of eggs produced in the hormone-treated groups was significantly higher than in the natural mating group, the rate of females that underwent copulation was unexpectedly low. Only 7 out of the 22 females were sperm positive in the superovulation group, while 11 out of 17 females in the natural mating group were sperm positive as determined by a vaginal plug or smear (Fig. S1). Thus, the fertilization rate in the hormone-treated groups was significantly lower than in the natural mating group (37 ± 15.2 % vs 89.5 ± 5.8 %; t-test, t_14_ = 3.83, p < 0.01). These results indicate that further optimization is required to outperform the natural mating procedure in producing zygotes or embryos for in vitro genome editing manipulation.

### Male presence during superovulation

To facilitate the receptivity of females after superovulation, males were introduced into female cages right after PMSG injection in some groups (#6, 7, 8), to allow females to become familiarized with the males. However, the timing of pairing during superovulation seemed to have no significant effect on the number of total or fertilized eggs between groups (t-test, t_20_ = 0.9, p = 0.38).

### Early development and in vitro culture of Nile grass rat embryos

To understand the time course of early embryo development and to determine permissive conditions and timing for embryo microinjection, eggs collected from three cohorts of superovulated females were cultured *in vitro*. None of the eggs collected from group #4 (19 hours post-hCG, Table 1) showed signs of fertilization at the time of harvesting; after 8 hours of culture in M2 medium, several zygotes developed pronuclei, but did not develop further (Fig. S2).

Previous studies have reported that in suboptimal culture conditions, early embryo development of mouse, hamster, and rat, arrests at the 1-or 2-cell stage, referred to as the ‘2-cell block’[46-49], which could be overcome with different concentrations of nutrients and culture media[50-53]. To bypass a potential 2-cell block in this species, 2-cell stage embryos (n = 11) collected on day 5 (52 hours post-hCG, group #3, Table 1) were cultured in M2 medium on a 37°C heat stage, in air for 5 hours. These culture conditions supported some embryos reaching the 4-cell stage. The embryos were then transferred either into Sydney IVF Fertilization Medium (SIFM) mouse embryo culture medium or modified rat 1-cell embryo culture medium (mR1ECM) with PVA, while a few were left in M2 medium. After 2 to 3 days of incubation at 37°C, 5% CO_2_, blastocysts were observed in both SIFM and mR1ECM media, but not in M2 medium (Fig. 1). Out of 5 embryos from each group, 4 developed into blastocysts in mR1ECM, while only 1 developed into blastocyst in SIFM; moreover, blastocysts appeared to be bigger with more cells in mR1ECM medium.

**Fig. 1.**
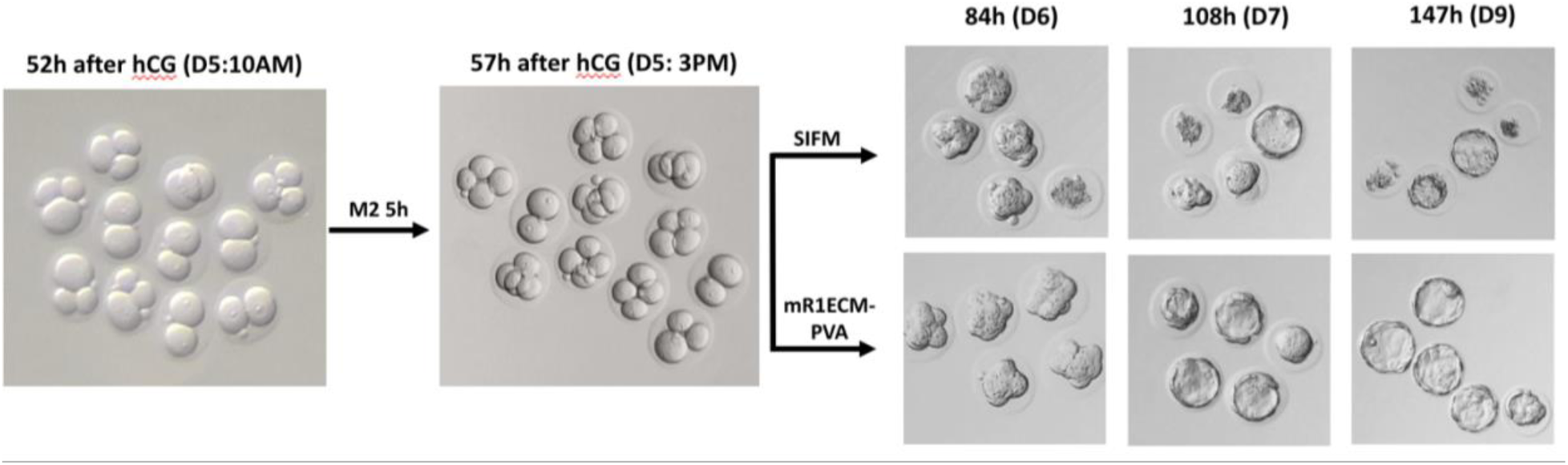
Comparison of two standard mouse and rat embryo media for grass rat embryos in vitro culture. Nile grass rat 2-cell stage embryos were flushed on the morning of day 5, 52 hours post hCG injection from oviducts of sperm-positive females. Embryos were washed in M2 medium and then cultured in mouse embryo culture medium SIFM, mouse embryo handling medium M2, or rat embryo culture medium mR1ECM-PVA. During in vitro culture, embryos were cultured in 20 µL medium micro-drops under mineral oil, in a 5% CO_2_ incubator at 37°C.

We further tested mR1ECM medium on 1-cell stage embryos to determine if a 2-cell block occurred. At the time of embryo collection from group #5 (27 hours post-hCG, Table 1), 4 out of 13 embryos appeared to have pronuclei; then, embryos were divided into 2 groups and cultured in mR1ECM media supplemented either with PVA or BSA. After 21 hours of culture, or 48 hours post-hCG, 5 out of 6 embryos in mR1ECM-PVA and 5 out of 7 in mR1ECM-BSA reached the 2-cell stage. Blastocysts started to appear at 96 hours post-hCG first in mR1ECM-BSA, and subsequently in mR1ECM-PVA (Fig. 2). Together these results demonstrate that mR1ECM media can support grass rat embryos to develop into blastocysts from the 1-cell stage *in vitro* and bypass a potential 2-cell block. The number of pronuclei, 4 out of 13 at 27 hours post-hCG, and the number of 2-cells, 10 out of 13 at 48 hours post-hCG, collectively indicated that most of the pronuclei developed 27-hours post-hCG, which would be coincide with the evening or night on day 4 post-mating. This suggests that manipulating grass rat embryos at the pronuclei stage would be inconvenient, unlike pronuclear microinjection of mice or rats that is routinely performed during the daytime, posing a practical challenge for gene editing of grass rats.

**Fig. 2.**
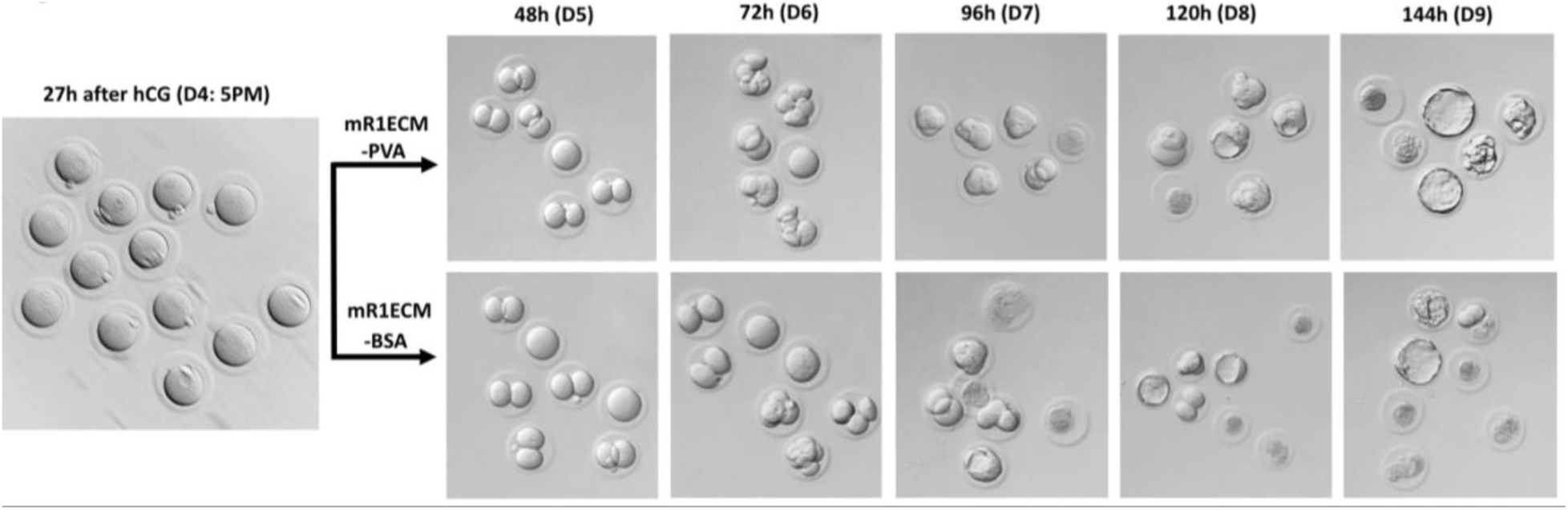
Nile grass rat embryos in vitro culture from 1-cell stage. Grass rat eggs were flushed on the afternoon of day 4, 27 hours post hCG injection from oviducts of sperm positive females. Eggs or zygotes were washed and then cultured in mR1ECM with PVA or with BSA, in a 5% CO_2_ incubator at 37°C until harvest.

### Rai1 gRNA validation and generation of Rai1 KO Nile grass rat via i-GONAD

Gurumurthy et al. developed GONAD and improved-GONAD (i-GONAD), in which CRISPR components are injected into the oviduct harboring fertilized eggs followed by electroporation allowing the delivery of CRISPR reagents into zygotes. i-GONAD does not require either embryo manipulation *in vitro*, or surrogate pseudopregnant females[39, 42]. The lack of established methods for production of pseudopregnant grass rat surrogates, pointed to i-GONAD as a viable approach to generate genome edited Nile grass rats.

The gene we targeted was *Rai1*, encoding a histone binding protein. Rai1 haploinsufficiency is responsible for Smith-Magenis Syndrome (SMS), a rare neurodevelopmental condition characterized by obesity, autistic behavior, and circadian rhythm and sleep disturbances[54, 55]. Although the obesity and behavioral traits have been well recapitulated in *Rai1*-knockout mice, *Rai1*^+/-^ mice ‘clearly differ from SMS patients’ regarding their sleep and circadian rhythms [56]. Contrary to the daytime sleepiness seen in SMS patients, the total time-spent-awake in *Rai1*^+/-^ mice was comparable to WT during their active phase. On the other hand, SMS patients also experience frequent night awakening, while *Rai1*^+/-^ mice slept significantly more than WT during their resting phase. These observations raise a possibility that the inverted chronotype contributes to the lack of sleep phenotypes in the SMS mouse model. Thus, *Rai1* is an ideal gene to test the utility of Nile grass rats for human disease modeling.

Two guide RNAs (gRNAs), g169 and g170, were designed to delete 2,035 bp of exon3, encoding most of the Rai1 protein (Fig. 3A). *In vivo* targeting efficiency of the gRNAs was validated in a female from natural mating that underwent i-GONAD. Within an hour following the i-GONAD procedure, three zygotes were retrieved from the oviduct and cultured in an incubator to the morula/blastocyst stage. After 4 days of culture in mR1ECM-PVA medium, embryos were collected and lysed individually for analysis by PCR amplifying both long-range and short-range amplicons of the target sites for g169 and g170. PCR analysis revealed that 1 out of the 3 embryos carried a large deletion between the cut sites of the two gRNAs (Fig. 3B & 3C). Subsequent sequencing data revealed that the embryo also carried an allele with indels at the cut sites of both g169 and g170, while another embryo carried a 6 bp mutation around g169 (Fig. 3D & 3E), indicating that 2 out of 3 embryos were successfully edited.

**Fig. 3.**
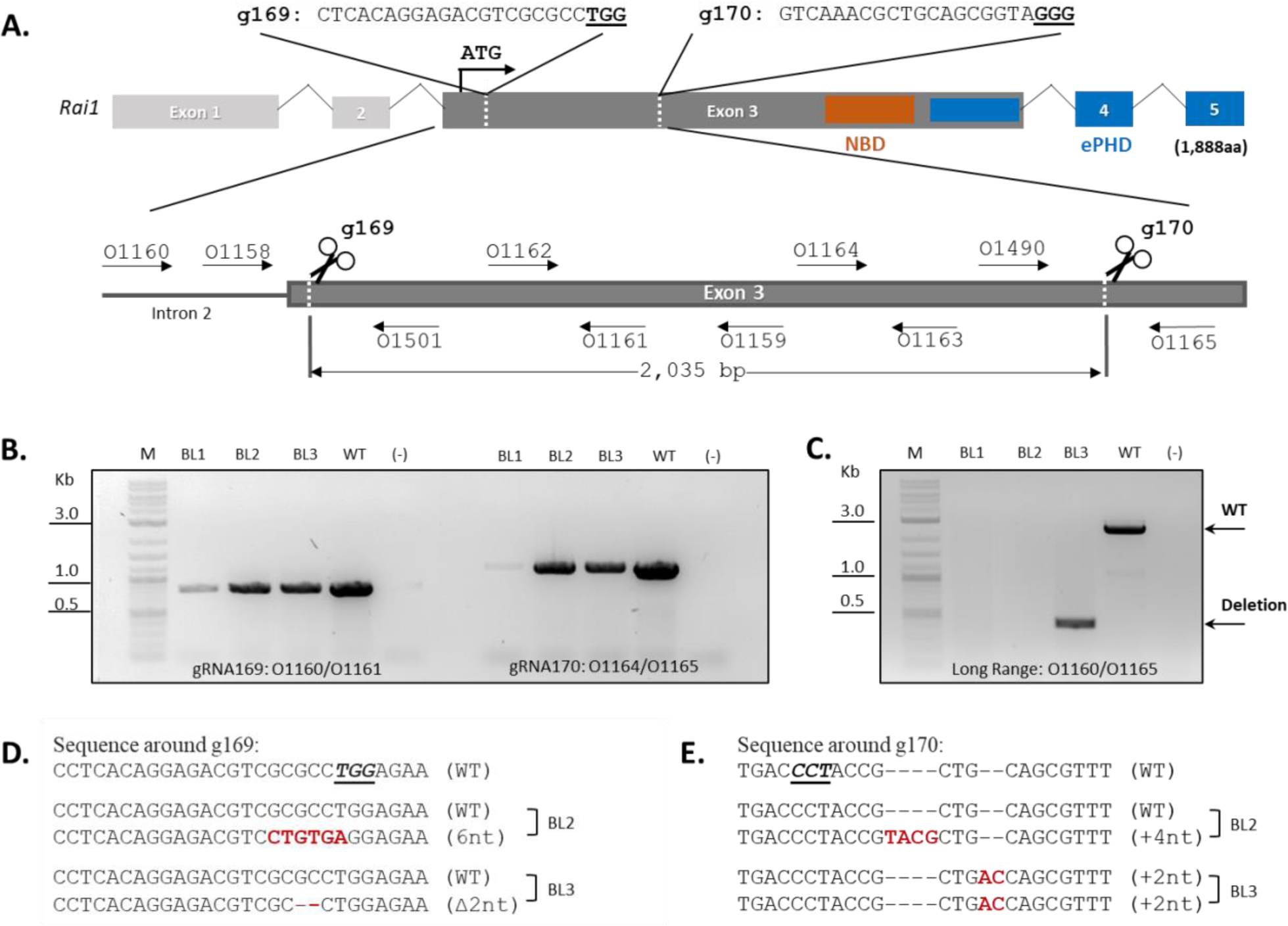
Targeting of *Rai1* in the Nile grass rat. A) A locus map denoting the location of gRNA cutting sites (dashed lines, PAM – underlined), primers (black arrows) relative to exons of the Nile grass rat *Rai1* gene. The locations of two predicted protein-interacting domains: a nucleosome-binding domain (NBD) and an extended plant homeo-domain (ePHD) are mapped to the corresponding coding region. B) PCR of the 2 target regions for blastocysts that underwent i-GONAD: no obvious difference between targeted sample and wild-type. C) Long range PCR spanning both target regions demonstrates that 1 out of 3 blastocysts carries a large deletion. D) and E) Alignments with reference genome demonstrate the presence of indel mutations around g169 and g170 target sites, not detectable by molecular weight differences in PCRs shown in B.

Once the editing efficiency of the g169 and g170 was confirmed, multiple cohorts of female grass rats, either superovulated with PMSG/hCG or following natural mating, underwent i-GONAD in attempts to generate *Rai1* knockout (KO) grass rats (Table 2). For the hormone-treated animals, PMSG was administered on day 1 at 6 am, hCG on day 3 at 2:00 pm, and i-GONAD was performed on day 4 at ∼5:00 pm following the confirmation of successful mating in the morning. For natural mating pairs, a vaginal smear was checked to confirm sperm presence in the morning of day 4 to day 5 post-pairing, and i-GONAD was performed in the afternoon.

**Table 2:** Outcome of Nile grass rat targeting and litters following i-GONAD.

| Group | Ovulation induction method | Male presence | # females on Day 1 | # i-GONAD females | # litters | # pups born | # edited pups |
| --- | --- | --- | --- | --- | --- | --- | --- |
| #1 | Superovulation | b/n hCG admin and vaginal smear | 32 | 10 | 0 | 0 | 0 |
| #2 | Postpartum natural mating | prior to i-GONAD | 16 | 8 | 0 | 0 | 0 |
| #3 | Superovulation | Day 1 - 10 | 13 | 2 | 1 | 5 <sup>1</sup> | 1 |
| #4 | Natural mating Phase 1 | Day 1 - 10 | 29 | 9 | 6 | 18 <sup>2</sup> | 0 |
| #5 | Natural mating Phase 2 | Day 1 - 10 | 20 | 12 | 8 | 25 | 2 |
| #6 | Natural mating Phase 3 | Day 1 - 10 | 8 | 4 | 2 | 9 | 4 |

<sup>1</sup>All pups died after birth. <sup>2</sup>Some pups died before biopsy.

Initially, females were singly housed after i-GONAD, but neither the PMGS/hCG primed (n = 10) nor those from natural mating pairs (n = 8) gave birth to any live pups (Table 2, group #1 & #2). Since previous studies of hamsters and voles have suggested that male proximity contributes to pregnancy success[57-60], in subsequent cohorts, the male was not removed from the cage after the i-GONAD procedure, and was co-housed with the female for at least 10 days to facilitate pregnancy maintenance. In the cohort of hormone-treated females (Table 2, group #3), 2 out of 13 were sperm positive as detected by vaginal smear and underwent i-GONAD. One of them produced a litter of 5 offspring, with one founder carrying a large deletion of *Rai1*, although the litter died a few days postnatally. On the other hand, 21 out of 49 females from natural mating pairs underwent i-GONAD, producing a total of 17 litters and at least 57 pups. From the first several litters, only 2 live pups out of 43 born carried *Rai1* large deletions (Fig. 4A). Subsequently, four females underwent i-GONAD and produced 2 litters of 9 pups in total (Table 2, group #6), of which 4 pups carried multiple large deletions (Fig. 4B). Sanger sequencing revealed various deletions across litters (Fig. 4B). The deletion events appeared to occur mostly heterozygously or exhibit mosaicism, except for animals iG5-iG8, which did not show WT bands (Fig. 4A and 4B). We reasoned that the absence of WT band may be due to inefficient PCR amplification of the larger WT amplicon, because another primer set amplifying smaller WT amplicon provided signal from animals iG6 and iG7 (Fig. 4C lower panel). In sum, analysis of founder (G0) offspring revealed successfully edited *Rai1* KO animals following i-GONAD delivery of CRISPR reagents.

**Fig. 4.**
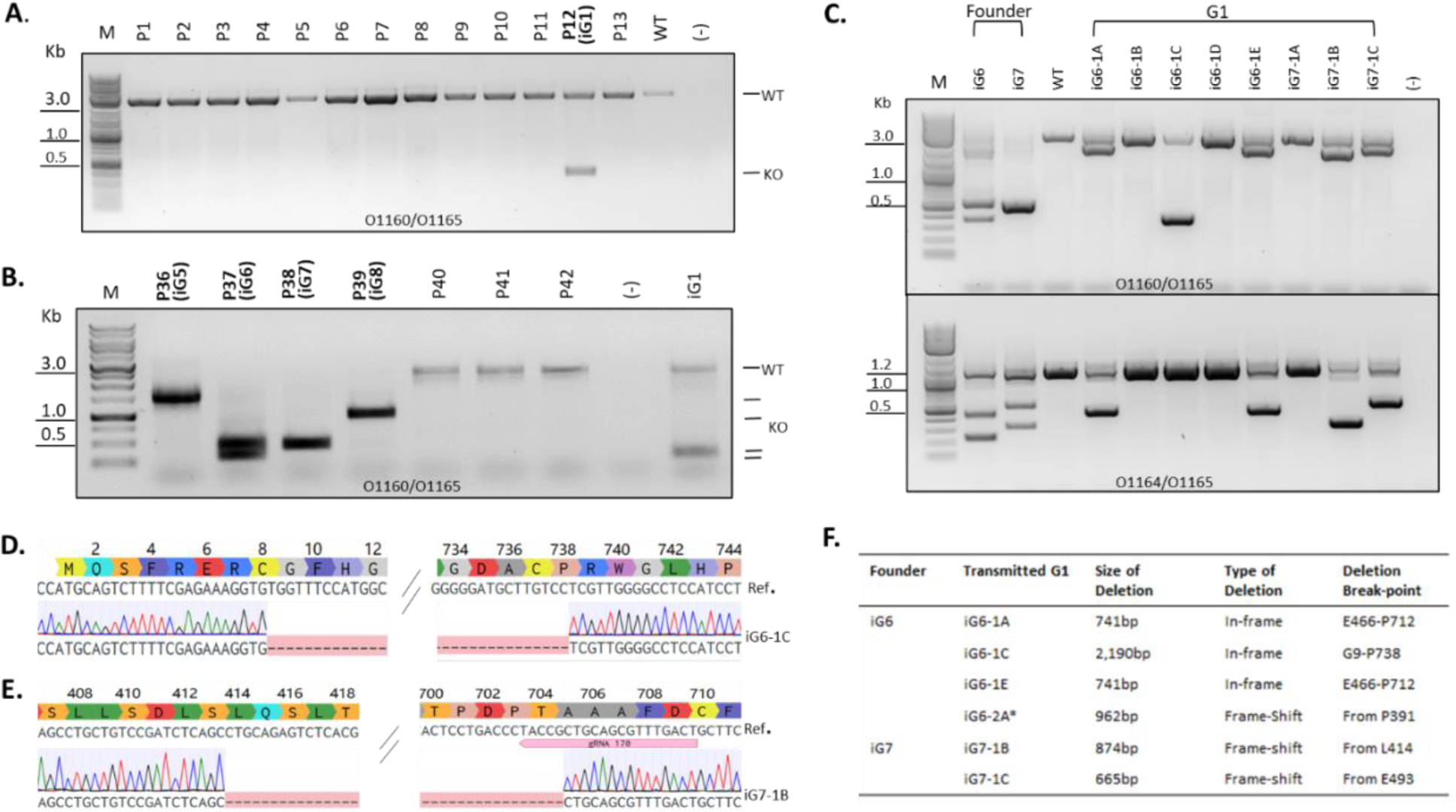
Generation of *Rai1*-edited Nile grass rat founders and G1 offspring via i-GONAD. A) Gel image of long-range PCRs of 13 pups (P1-P13) from the first 4 i-GONAD litters. Founder P12 (iG1) carries a large deletion. B) Gel image of long-range PCRs from a representative litter (pups P36 - P42), which produced 4 large deletions out of 7 pups. C) PCRs demonstrating that multiple deletions from 2 founders, iG6 and iG7 were transmitted to G1 offspring. D) Sanger sequencing chromatogram of G1 animal iG6-1C from founder iG6 shows a 2,190bp deletion. E) Sanger sequencing chromatogram of iG7-1B, one of the G1 offspring from founder iG7 shows transmission of a 874bp deletion. F) Summary of transmitted deletion alleles from founders iG6 and iG7. *This G1 animal was from a second litter of founder iG6 shown in Fig. S3.

After breeding with WT animals, *Rai1* KO founders iG6 and iG7 successfully transmitted the edited *Rai1* allele to their G1 offspring (Fig. 4C). Founder iG6 transmitted 2 in-frame deletions to G1: a 2,190 bp deletion removing amino acids G9-P738 (Fig. 4D), and a 741 bp deletion of amino acids E466-P712. In a subsequent litter from iG6, a frameshift deletion of 962 bp was also transmitted to G1 (Fig. S3). Founder iG7 transmitted 2 frameshift deletions, 874 bp (Fig. 4E) and 665 bp were detected in G1 animals iG7-1B and iG7-1C (Fig. 4F). Multiple bands were detected in individual founders, indicative of mosaicism – the presence of multiple alleles in the same animal, whereas only single altered DNA species were detected in G1 animals (Fig. 4C and 4D). These results demonstrate successful gene targeting of *Rai1* in Nile grass rats and stable transmission of the targeted allele to the next generation.

### In vitro Nile grass rat embryo microinjection

i-GONAD is suitable for delivery of CRISPR RNPs, mRNA, gRNA and ssODN. However, the delivery of large DNA molecules harboring transgenes requires microinjection directly into the zygote despite emerging reports about introducing large DNAs in vivo by adeno-associated virus[61-63]. Thus, being able to deliver genetic material via microinjection is a key step towards sophisticated genetic manipulations such as conditional gene targeting. To this end, we tested microinjection of *Rai1* CRISPR reagents directly into early embryos *in vitro*. Seven 2-cell and three 4-cell stage embryos were collected on day 5 following natural mating and subjected to microinjection of the same *Rai1* CRISPR reagents. HEPES buffered mR1ECM-BSA medium was used for embryo collection and microinjection of RNPs, while mR1ECM-PVA was used for embryo culture post-microinjection. During the microinjection process, none of the embryos showed any sign of cytotoxicity or morphological changes. Following 3 days of culture in mR1ECM-PVA medium, 9 out of 10 embryos developed into 5 blastocysts and 4 morulae. PCR and sequencing of the amplicon spanning the region between the two *Rai1* gRNAs revealed that 7 out of 9 embryos carried large deletions of the *Rai1* gene (Fig. 5). Therefore, CRISPR RNP microinjection into grass rat embryos can result in high-efficiency genome editing.

**Fig. 5.**
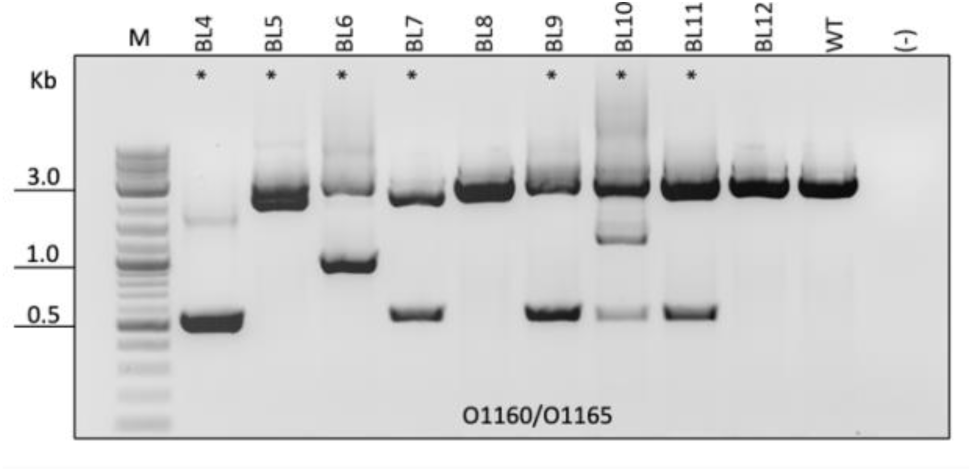
Targeting efficiency of *Rai1* gRNAs by 2-cell microinjection of in vitro cultured embryos. Gel image showing that 7 out of 9 embryos carry deletions with bands at lower molecular weights than WT.

## Discussion

CRISPR-Cas-based genome editing is not only used routinely in creating standard laboratory rodent models like mice and rats, but has also been used in engineering of non-conventional rodent models, including prairie voles[64, 65] and hamsters[66, 67], as well as livestock[30, 31, 34, 68]. However, genome editing has faced some unique challenges in diurnal rodents. Previous attempts to generate a germline transgenic line using a closely related diurnal rodent, the Sudanian grass rat (*Arvicanthis ansorgei*), reported repeated failures, likely due to “the lack of knowledge of experimental procedures suitable for creating transgenic diurnal rodents”[69]. The present study serves as the first step toward developing the diurnal Nile grass rat as a genetically tractable model for translational research. Through this effort, we have accomplished a few major milestones, including establishing conditions for superovulation, fertilization, and embryo culture, followed by i-GONAD to successfully produce founder animals carrying targeted deletions that were then transmitted to G1 offspring.

An effective protocol to generate enough embryos is critical for successful genome editing. Extensive reproductive biology research has established superovulation, *in vitro* culture, and *in vivo* fertilization protocols in rodent species including mice, rats, hamsters[70] and prairie voles[71], laying the foundation for successful genome editing in those species, but such reproductive and early embryo development studies are lacking for the Nile grass rat or other diurnal rodents. The results from the current study contribute a working protocol that can effectively produce a large number of eggs in grass rats. Although the rate of females showing signs of successful mating was lower in the superovulated group than in the natural mating group, the ability to produce more eggs will be useful for *in vitro* fertilization approaches which are advantageous for reducing the number of egg donors to be euthanized while obtaining large number of embryos [72, 73]. Thus, this technique could be used to assist future genome editing in grass rats or other diurnal rodents.

Conditions that support early embryo development *in vitro* enable embryo manipulation required for delivery of genome editing reagents such as microinjection and electroporation as well as the study of early embryogenesis. Through our superovulation studies, we were able to get a glimpse of the early embryo development timeline in the grass rats. Currently, there are no reports of *in vitro* handling of grass rat embryos or the timing and staging of early grass rat development. Based on the stage of embryos collected at different intervals from 19 to 52 hours post hCG injection, the time course of the early grass rat development could be mapped out as follows: fertilization completes < 19 hours; pronuclei form ≥ 27 hours; then 2-cells form > 40 hours. We found that both M2 and mR1ECM-HEPES could be used as short-term embryo handling medium, while mR1ECM-PVA and mR1ECM-BSA both support grass rat embryos to develop from the pronuclear to blastocyst stage in culture *in vitro*. These results suggest that media optimized for rats might be suitable for the Nile grass rat for further studies such as *in vitro* fertilization or embryo or sperm cryopreservation.

Understanding the timeline of natural mating, from breeding pair setup, vaginal plug and sperm detection to embryo harvesting and early embryo development, ultimately enabled us to perform genome editing in this species without superovulation. Furthermore, our initial attempts of i-GONAD with 18 females, which failed to produce any pups, led us to discover another unique feature of the reproductive success in this species - the requirement of male presence in order to carry pregnancies to term. While it is standard practice to single house mice or rats following embryo implantation or i-GONAD procedures, male proximity seems to be critical for successful pregnancy in grass rats. Similarly, it has been reported that for prairie voles continued males presence facilitates pregnancy maintenance[74]. Hence, the final piece of the puzzle was in place for targeting grass rats with the outcome of 5 founders surviving to adulthood. So far, 2 of the *Rai1* knockout founders have transmitted their deletion to G1 offspring after backcrossing with wild-type animals, demonstrating that the targeted alleles could be established as stable genome-edited grass rat lines for future functional studies.

## Conclusion

In the present study, fundamental steps were taken towards creating genome edited diurnal rodent models. The newly created *Rai1* KO Nile grass rat line using i-GONAD is a unique model for understanding the role of *Rai1* in the neurodevelopmental disorder SMS. More importantly, the high targeting rate of 2-cell embryo microinjection demonstrated its potential for other forms of gene editing, including the generation of point mutations, knocking in epitope tags and larger insertions, and creating conditional alleles with the Cre-loxP system. To facilitate future genome targeting in this species, we propose a scheme for Nile grass rat genome targeting, either through natural mating or via superovulation (Fig. 6). Briefly, if the day of pairing females with males in natural mating, or the day of PMSG administration is defined as day 1 (D1), females with a sperm positive vaginal smear on day 4 (D4) will be suitable for embryo targeting on day 5 (D5). Late evening of D4 or daytime of D5 is the embryo manipulation window for pronuclear or 2-cell staged embryos. Co-housing a female that underwent targeting with a male until at least day 10 (D10) is critical for the maintenance of a successful pregnancy. We hope this method will help guide future development of genetically modified grass rats and other diurnal rodents, which will promote greater utility of these models in basic and translational research.

**Fig. 6.**
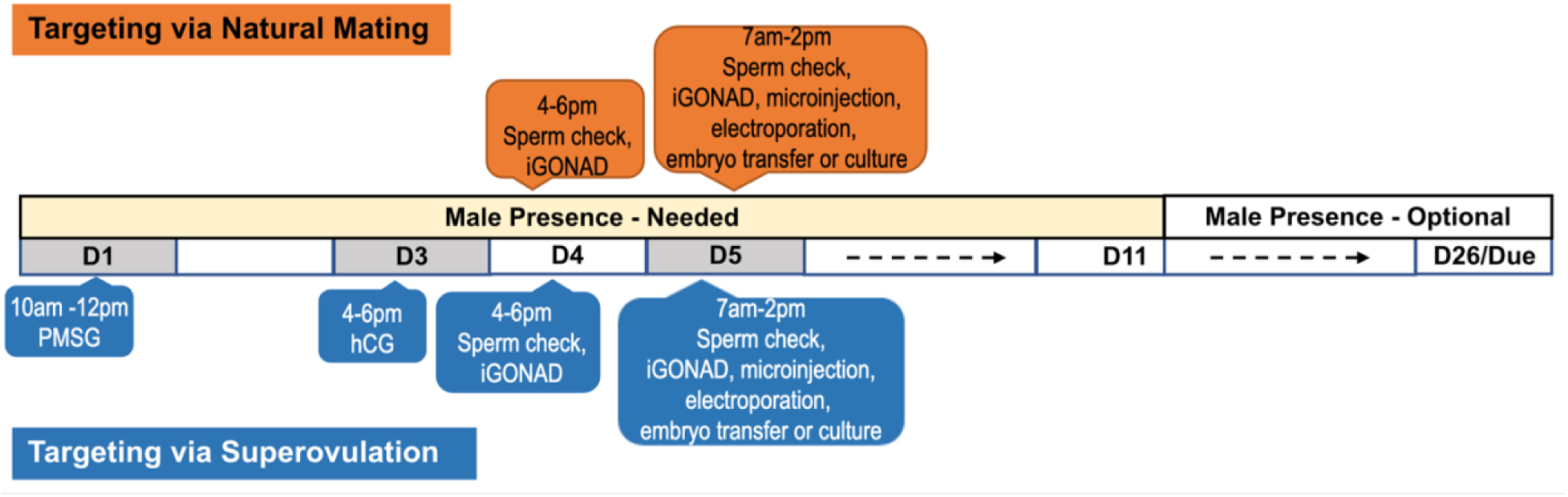
A proposed scheme for Nile grass rat genome targeting. Females ranging from 3 to 11 months old can be used as egg donors or embryo transfer recipients, while proven breeder males are needed for mating. Procedure for natural mating: 1) The day that males are paired with females is defined as day 1 (D1). 2) Vaginal smear cytology is assessed in the late afternoon of day 4 (D4) to determine if i-GONAD could be performed. At that timepoint, zygotes were found to be either at or prior to the early pronuclear stage, so in vitro manipulation on the afternoon of day 4 is not recommended. 3) Vaginal smear cytology is assessed on the morning of day 5 (D5), to determine if embryo manipulation can be performed on that day, including i-GONAD, *in vitro* embryo electroporation, or microinjection followed by embryo transfer into surrogate females, or embryo culture *in vitro* for gRNA validation. 4) Females are co-housed with males at least until day 10 (D10). 5) Females are monitored for signs of labor starting from day 26 through day 30. 6) Any pups born are genotyped upon weaning.

## Methods

### Animals

Adult male and female Nile grass rats were obtained from an in-house laboratory colony at Michigan State University [15]. The colony was maintained in standard animal housing room under a 12:12 hour light/dark (LD) cycle, with lights on at 6:00 and lights off at 18:00. A metal hut was provided in each cage for shelter and enrichment. Food (Prolab 2000 #5P06, PMI Nutrition LLC, MO, USA) and water were available *ad libitum.* All procedures were conducted in accordance with the National Institutes of Health Guide for the Care and Use of Laboratory Animals (NIH Publication No. 80-23) and were approved by the Institutional Animal Care and Use Committee of Michigan State University.

### Superovulation and natural mating

As a species of induced ovulators, female grass rats only start the ovulatory cycle after being co-housed with a male. Animals reach reproductive maturity around 2-months of age and can reproduce up to 16-months of age based on our observation of colony animals. Young to middle-aged adults (3-to 7-month-old) were used in this experiment based on the availability of animals in the colony. For superovulation experiments, singly housed females were first injected with PMSG (15 IU or 20 IU, BioVendor) in the morning (time of injection and dosage are listed in Table 2), followed by hCG (CHORULON®, MERCK) at various intervals ranging from 36 to 57 hours, at the same dosage as PMSG. A male was introduced to each hormone treated female to allow mating, either following hCG administration or immediately after injection of PMSG. For natural mating, females were paired with males at the ratio of 1:1. In the initial experiments a vaginal smear from each female under mating was checked daily, until a plug or sperm was found. In later experiments, vaginal smear and sperm presence were checked from day 4 to day 5 post pairing.

### Embryo collection and culture

Females that had successfully mated, as confirmed by a vaginal plug or sperm positive smear, were euthanized with sodium pentobarbital (i.p. 150 mg/kg). Bilateral oviducts were dissected out and placed in HEPES-buffered embryo culture medium either 27- or 56-hours post hCG injection. Embryos were released either by tearing the ampulla at the day of plug, or oviduct flushing the next day after plug or presence of sperm. After washing in HEPES-buffered M2 medium (Sigma) or mR1ECM-BSA (CytoSpring LLC), embryos were cultured in either M2 medium (Sigma), mouse embryo culture medium SIFM (COOK Medical), or rat embryo culture media mR1ECM supplemented with PVA or BSA (CytoSpring LLC).

### Reagents for genome targeting

CRISPR-Cas9 technology was used as a genome editing tool. Two guide RNAs (gRNAs) targeting the *Rai1* (Gene ID: 117711603, https://www.ncbi.nlm.nih.gov/gene/117711603/; Gene: ENSANLG00005011239, <u>Gene: RAI1 (ENSANLG00005011239) - Summary -</u> <u>Arvicanthis_niloticus_GCA_011762505.1 - Ensembl 108</u>) were designed using Benchling (Benchling [Biology Software] 2021) and synthesized as single gRNA by IDT (Integrated DNA Technologies). Both gRNAs targeted exon 3 of *Rai1*, with protospacer and PAM sequences 5’-CTCACAGGAGACGTCGCGCC - TGG 3’ (g169) and 5’– AGTCAAACGCTGCAGCGGTA – GGG 3’ (g170). Ribonucleoprotein (RNP) complexes were assembled in vitro by incubating gRNAs with wild-type S.p. Cas9 Nuclease 3NLS protein (IDT) at 37°C for 5 minutes, and then kept on ice. For i-GONAD, 1 µL Trypan blue solution (0.4%) and Duplex buffer (IDT) were added to the RNP mix to a final concentration of 200 ng/µL for each RNP.

### In vivo Genome Editing using i-GONAD

The i-GONAD procedure was performed as described previously[45]. Briefly, in the afternoon at the day of plug or sperm found, females were placed under isoflurane anesthesia, oviducts would be exposed as in a standard embryo transfer procedure, after the RNP mixture was delivered into oviducts with a glass pipette, 4 pulses of 50 V were delivered using a pair of disk electrodes connected to the electroporator, Genome Editor (BEX CO., LTD). After i-GONAD surgeries were completed, females were placed back into their home cage, with or without a male, and were monitored daily.

### In vitro Genome editing

Genome editing was conducted *in vitro* via microinjection. RNPs were diluted with 10mM Tris pH7.5 buffer to a final concentration of 50 ng/µL each RNP. Embryo donor females were euthanized the next morning after vaginal sperm presence was confirmed. Embryos were collected and placed in HEPES-buffered mR1ECM-BSA culture medium, 2-cell nuclear microinjection was performed using CELLectro (a gift from Dr. Leyi Li, Cold Spring Harbor Lab), as previously described [75, 76]. Microinjected embryos were cultured in an incubator (5% CO_2_; 37°C), until later stages. Morulae or blastocysts were collected and genotyped individually to confirm CRISPR editing efficiency *in vitro* using PCR.

### PCR Genotyping

A small number of targeted embryos were genotyped to evaluate the editing efficiency of gRNAs. Briefly, embryos cultured in vitro were harvested when they were at blastocyst or morula stage, after 4 days in culture. Each embryo was placed individually into a PCR tube containing 10 µL of tail lysis buffer (0.1 mg/mL Proteinase K in 0.5% Triton X-100, 10mM Tris pH8.5). Before they could be used as genomic DNA templates in PCR reaction, the embryo lysis would go through two steps: digestion at 56°C for at least 1 hour and heat treatment at 85°C for 15 min.

Pups born from females that went through i-GONAD or embryo transfer were genotyped using standard procedures. In brief, small ear biopsies were lysed at 56°C overnight for PCR using tail lysis buffer described above, then heat treated at 85°C for 15 min. Sanger sequencing (GENEWIZ Inc., and Quintara BioSciences) was performed on purified PCR amplicon DNA. Primers used for sequencing and genotyping are listed in Table 3.

**Table 3.**
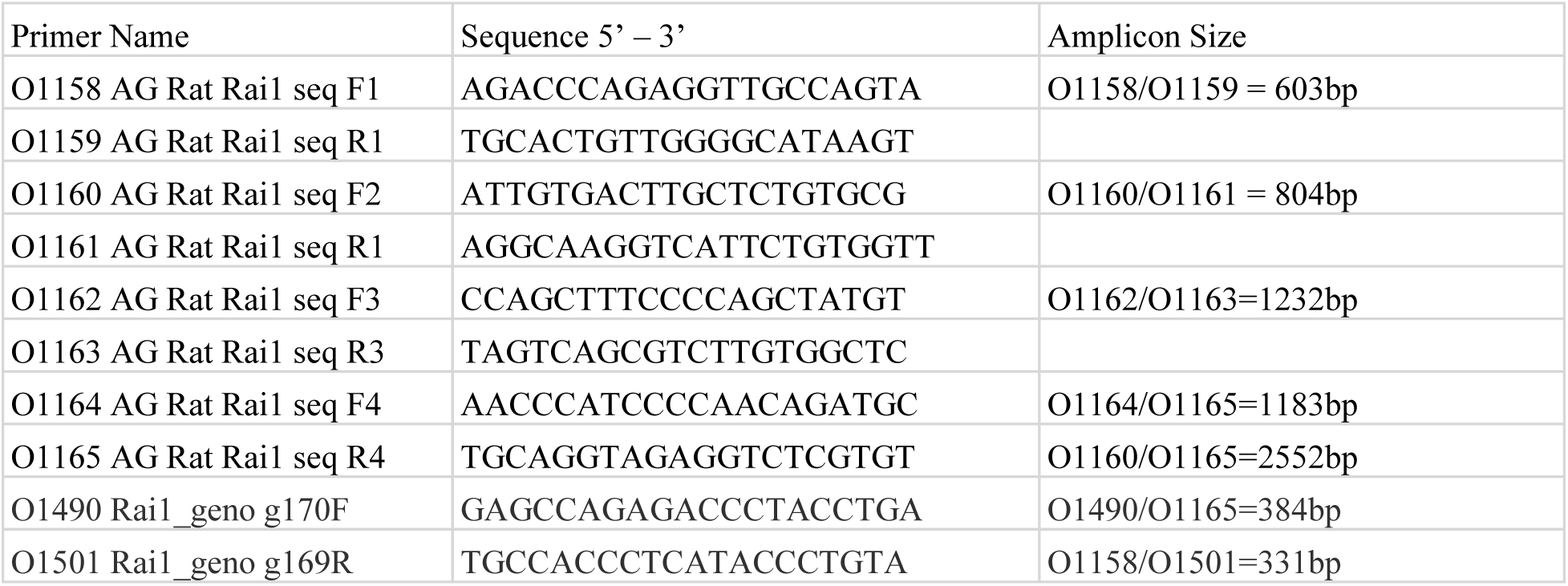
Primers for *Rai1* genotyping.

**Fig. S1.**
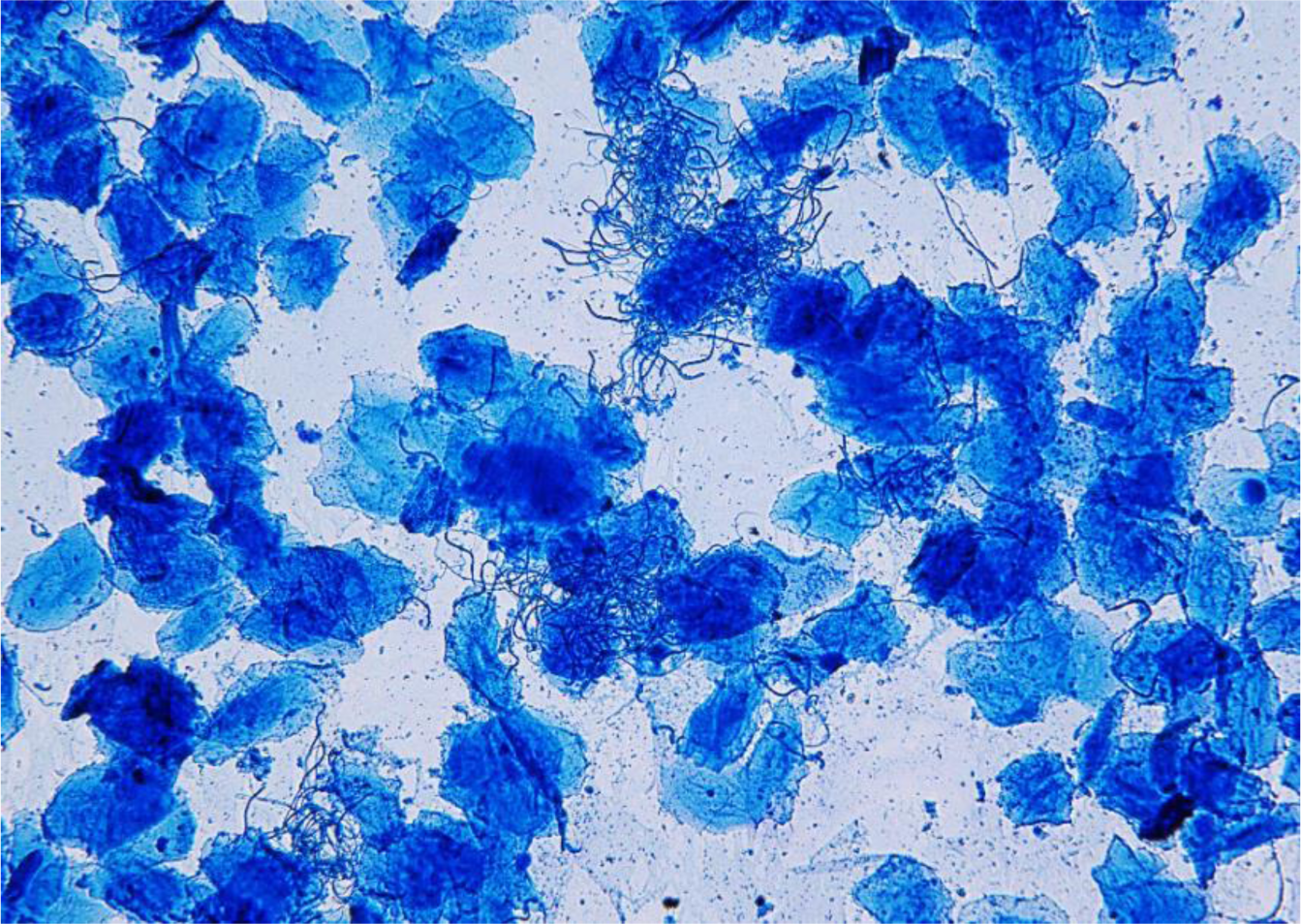
Representative sperm-positive vaginal smear cytology of a female Nile grass rat. Sperm with a thin, elongated, hair-like appearance, and cornified epithelial cells, larger globular objects, are stained with methylene blue.

**Fig. S2.**
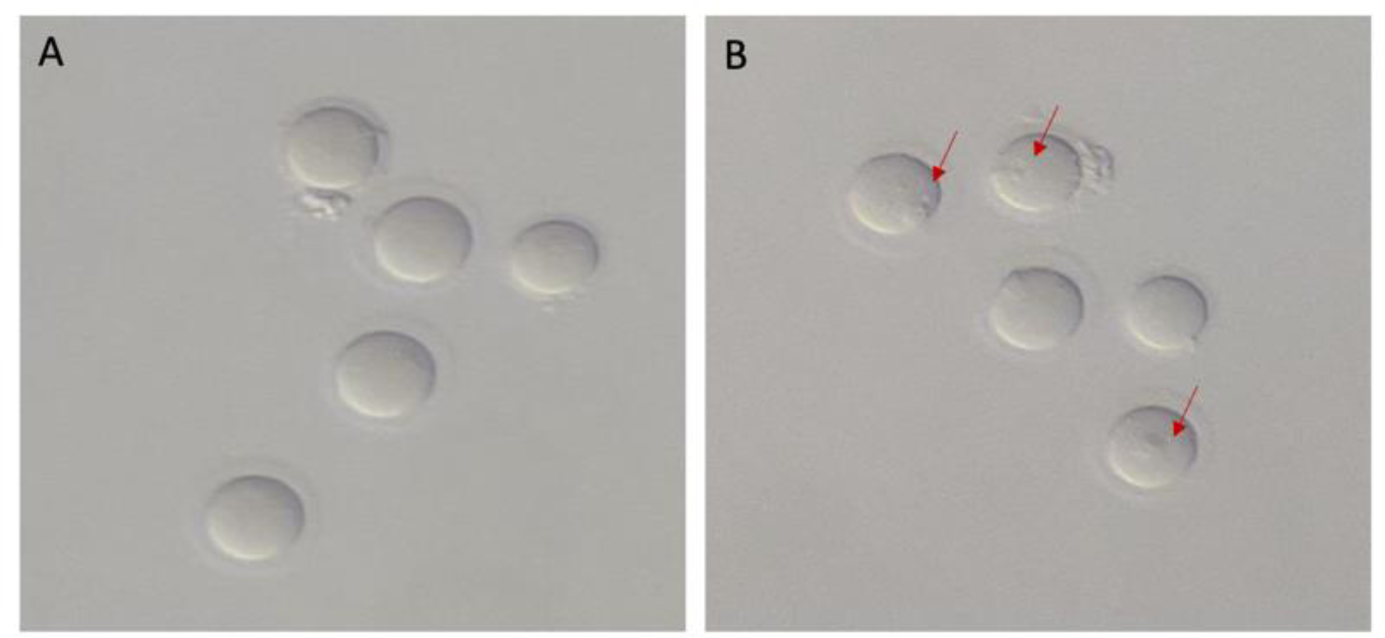
Eggs retrieved from sperm-positive female grass rats the day after hCG administration and mating. A) No pronuclei are visible in eggs collected 19 hours after hCG. B) Pronuclei (red arrows) are present in zygotes cultured to 27 hours after hCG.

**Fig. S3.**
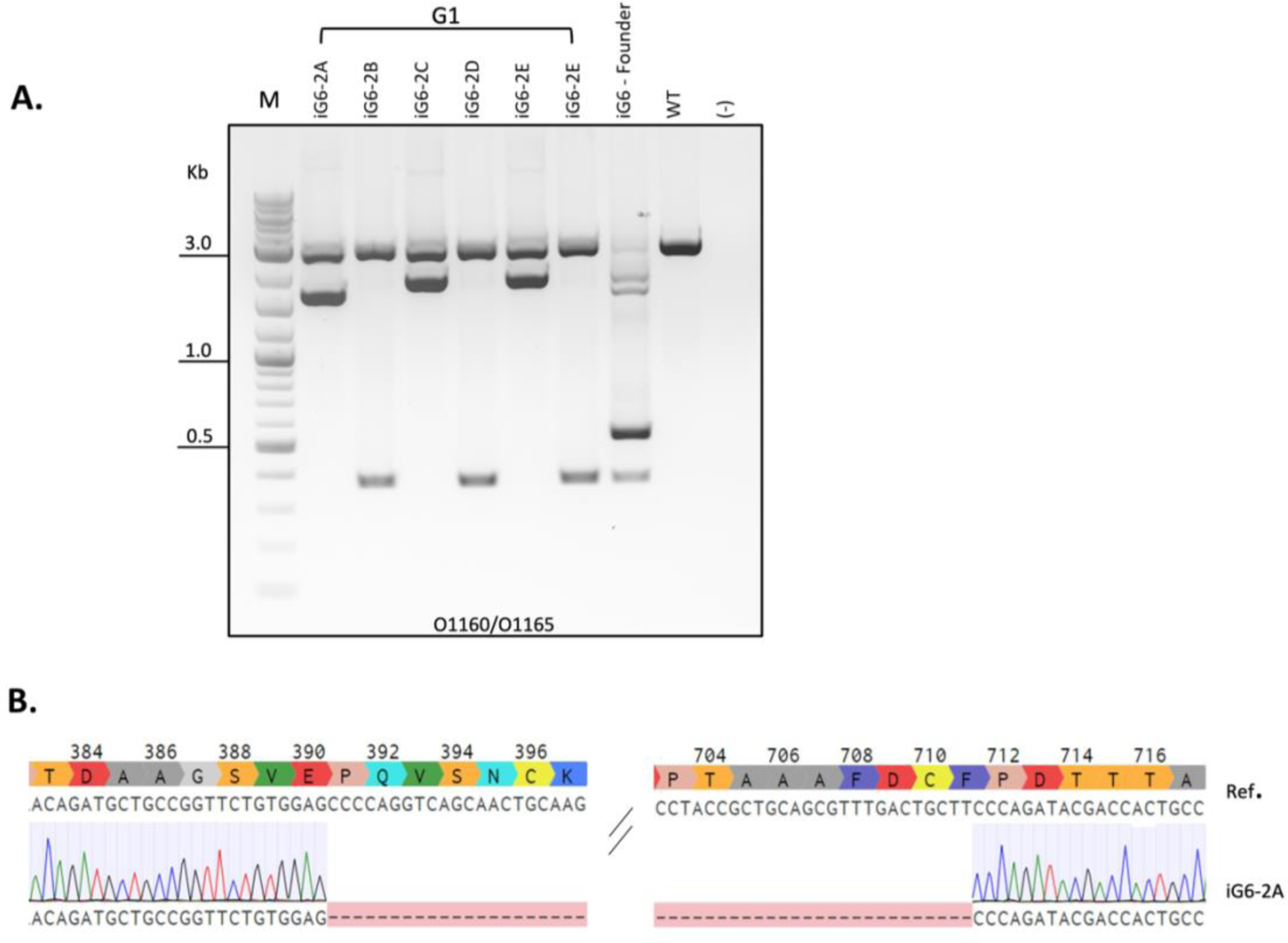
Additional G1 transmission of Nile grass rat *Rai1* deletion. A) Gel image showing long range PCR of G1 offspring from founder iG6 mated with a WT animal. Three different deletions were present in 6 pups assessed. B) Sanger sequence chromatogram of G1 animal iG6-2A shows a 962bp deletion, which results in frameshift after P391.

## Declarations

### Ethics approval and consent to participate

All procedures were conducted in accordance with the National Institutes of Health Guide for the Care and Use of Laboratory Animals (NIH Publication No. 80-23) and were approved by the Institutional Animal Care and Use Committees of Michigan State University and University of California Santa Barbara.

### Consent for publication

The authors give their consent for this manuscript to be published in the journal Genome Biology.

### Availability of data and materials

all data are presented in the manuscript.

### Competing interest

The authors declare that they have no known competing interests that could have appeared to influence the work reported in this paper.

### Funding

The work is supported by a MSU College of Social Science Faculty Initiative Fund to LY and NIH/NINDS R21NS125449 to SI and LY. HX, EYD and BA are supported by the MSU Global Impact Initiative Funds. BZ and SI are supported by the Jim and Sandy Danto Family Research Fund.

## Authorship Contributions

HX: Designed the study, performed embryo culture and manipulation experiments, created *Rai1*

KO grass rats, performed data analysis, initial draft with LY, revised manuscript.

KL: Performed animal husbandry, performed superovulation experiments, assisted in the creation of *Rai1* KO grass rats.

EYD: Contributed to the target design, revised manuscript.

HT: Contributed to the design of superovulation experiment, revised manuscript.

BA: Genotyped the *Rai1* KO grass rats.

JS: Assisted in the creation of *Rai1* KO grass rats and superovulation experiments.

BZ: Assisted in the annotation of *Rai1* gene and protein in grass rats.

SI: Conceived the project, acquired funding, revised manuscript.

LY: Conceived the project, acquired funding, assisted in the creation of *Rai1* KO grass rats, performed data analysis, initial draft with HX, revised manuscript.

## Acknowledgements

We thank Dr. Leyi Li, Cold Spring Harbor Laboratory, for providing the CELLectro microinjection device.

## Notes

### Competing Interest Statement

The authors have declared no competing interest.

